# Representation of real-world event schemas during narrative perception

**DOI:** 10.1101/252718

**Authors:** Christopher Baldassano, Uri Hasson, Kenneth A. Norman

**Author notes:** **Corresponding author**: Christopher Baldassano, Columbia University, 1190 Amsterdam Ave., New York, NY 10027. **Conflict of Interest**: “The authors declare no competing financial interests.”.

## Abstract

Understanding movies and stories requires maintaining a high-level situation model that abstracts away from perceptual details to describe the location, characters, actions, and causal relationships of the currently unfolding event. These models are built not only from information present in the current narrative, but also from prior knowledge about schematic event scripts, which describe typical event sequences encountered throughout a lifetime. We analyzed fMRI data from 44 human subjects presented with sixteen three-minute stories, consisting of four schematic events drawn from two different scripts (eating at a restaurant or going through the airport). Aside from this shared script structure, the stories varied widely in terms of their characters and storylines, and were presented in two highly dissimilar formats (audiovisual clips or spoken narration). One group was presented with the stories in an intact temporal sequence, while a separate control group was presented with the same events in scrambled order. Regions including the posterior medial cortex, medial prefrontal cortex (mPFC), and superior frontal gyrus exhibited schematic event patterns that generalized across stories, subjects, and modalities. Patterns in mPFC were also sensitive to overall script structure, with temporally scrambled events evoking weaker schematic representations. Using a Hidden Markov Model, patterns in these regions can predict the script (restaurant vs. airport) of unlabeled data with high accuracy, and can be used to temporally align multiple stories with a shared script. These results extend work on the perception of controlled, artificial schemas in human and animal experiments to naturalistic perception of complex narrative stimuli.

**Significance Statement:** In almost all situations we encounter in our daily lives, we are able to draw on our schematic knowledge about what typically happens in the world to better perceive and mentally represent our ongoing experiences. In contrast to previous studies that investigated schematic cognition using simple, artificial associations, we measured brain activity from subjects watching movies and listening to stories depicting restaurant or airport experiences. Our results reveal a network of brain regions that is sensitive to the shared temporal structure of these naturalistic situations. These regions abstract away from the particular details of each story, activating a representation of the general type of situation being perceived.

## Introduction

Everyday perception involves processing and reacting to a rich, rapidly changing stream of sensory information. A large body of work in cognitive psychology has shown that “common sense” comprehension requires not just bottom-up feature recognition, but also activation of knowledge schemas about the expected structure of the world (Bartlett, 1932; Piaget, 1926; Zacks, Speer, Swallow, Braver, & Reynolds, 2007).

Numerous recent studies, in both humans and animals, have explored how schemas are stored in the brain and how they influence ongoing processing. A critical region implicated in many of these studies is the medial prefrontal cortex (mPFC). This region shows encoding-related activity predictive of subsequent memory for schema-congruent stimuli (van Kesteren et al., 2013), increased activity when remembering schematic knowledge (Brod, Lindenberger, Werkle-Bergner, & Shing, 2015), and up-regulation of intermediate early gene expression when assimilating new information into a schema (Tse et al., 2011), and damage to the mPFC results in deficits for schema-related processing (Ghosh, Moscovitch, Melo Colella, & Gilboa, 2014). The mPFC is thought to mediate schematic activation in a network of regions, including the hippocampus (van Kesteren, Fernández, Norris, & Hermans, 2010; van Kesteren, Ruiter, Fernández, & Henson, 2012) and cortical regions such as posterior cingulate and angular gyrus (Gilboa & Marlatte, 2017).

However, the connection between this work and real-world perception has been lacking. The vast majority of these neuroscientific studies of schema have used simple, arbitrary relationships such as flavor-place associations (Tse et al., 2007, 2011) or associations between artificial stimuli (Brod, Lindenberger, & Shing, 2016; Brod et al., 2015). In naturalistic situations, we make use of highly elaborated schemas that have been learned and consolidated throughout our lifetimes, which may or may not rely on the same neural mechanisms as novel schemas involving simple relational binding. Also, most of these studies (Brod et al., 2015; van Kesteren et al., 2010) only compare schematic versus non-schematic conditions, making it unclear whether these regions actually represent information about *which* schema is active, or are engaged when *any* schema is active (regardless of content).

In this study we examine a specific type of naturalistic temporal schema, called a “script.” Schank & Abelson (1977) described a script as “a predetermined, stereotyped sequence of actions that defines a well-known situation,” and proposed that “most of understanding is script-based.” This central role of scripts was echoed by Mandler (1984), who proposed that scripts “are intimately involved in most of our daily processing, and an understanding of their structure and how it is used would add materially to our understanding of how the mind works.” These theoretical proposals were followed by empirical studies of the contents and consistency of scripts across people, which found that subjects largely agreed on how to segment activities into events as well as the typical “characters, props, actions, and the order of the actions” (Bower, Black, & Turner, 1979).

Using complex movies and audio narratives, we presented subjects with stories conforming to two different scripts that are highly familiar to our subject population: eating at a restaurant, and going through the airport. The stories within each script all shared the same high-level sequence of events (e.g. entering the restaurant, being seated, ordering, and eating food) but were highly dissimilar in terms of their low-level features (spoken words vs audiovisual movies), genres (e.g. science fiction vs thriller), characters, emotional content, and relative lengths of each of the constituent events. Using multiple converging analyses, we found that default mode regions, especially posterior medial cortex (PMC), mPFC, and superior frontal gyrus (SFG), exhibited sequences of activity patterns that were specific to each of the two scripts. These schema-specific event representations generalized across stories, across subjects, and across modalities, and were robust enough to be detected in held-out stories even without manual labeling of the events. Additionally, a separate control experiment showed that presenting events in scrambled order disrupted schematic effects in mPFC, providing evidence that this region is sensitive to the overall temporal structure of familiar scripts.

## Materials and Methods

### Subject details

We collected data from a total of 45 subjects (22 female, ages 18-38), 32 for the main experiment and 13 for the control experiment (described below). Subjects were native English speakers, in good health and with normal or corrected-to-normal vision. The experimental protocol was approved by the Institutional Review Board of Princeton University, and all subjects gave their written informed consent. To detect outlier subjects, each subject’s average posterior medial timecourse cortex (as defined below) across all stimuli was correlated with the mean timecourse of all other subjects (within the same experiment), to ensure that the subject had attended to and understood the narrative (Stephens, Silbert, & Hasson, 2010). One subject in the main experiment was excluded due to a correlation value that was more than 2.5 standard deviations below the rest of the group (r<0.15).

### Stimuli

The stimuli were designed to conform to two naturalistic schematic scripts that we expected to be familiar to all our subjects: eating at a restaurant, or catching a flight at an airport. Each scenario consisted of four events. For the restaurant stories, the events were: entering and being taken to a table, sitting with menus, ordering food and waiting for its arrival, and food arriving and being eaten. For the airport stories, the events were: entering the airport, going through the security checkpoint, walking to and waiting at the gate, and getting onboard the airplane and sitting in a seat.

Each story was approximately 3 minutes long (Figure 1). To identify schema representations that were modality-invariant, we presented 4 audio-visual movies and 4 spoken narratives for each of the two schemas. The stories all involved different characters and spanned multiple genres, sharing only the same high-level schematic script. The movies were sampled from films in which the restaurant schema was depicted *(Brazil, Derek, Mr. Bean, Pulp Fiction)* or the airport schema was depicted *(Due Date, Good Luck Chuck, Knight and Day, Non-stop),* and were edited for time and to conform as closely as possible to the four-stage schema script. The audio narratives were adapted from film scripts with a restaurant scene *(The Big Bang Theory, The Santa Clause, Shame, My Cousin Vinny)* or an airport scene *(Friends, How I Met Your Mother, Seinfeld, Up in the Air),* also edited for length and to match the schematic script. All narratives were read by the same professional actor.

**Figure 1:**
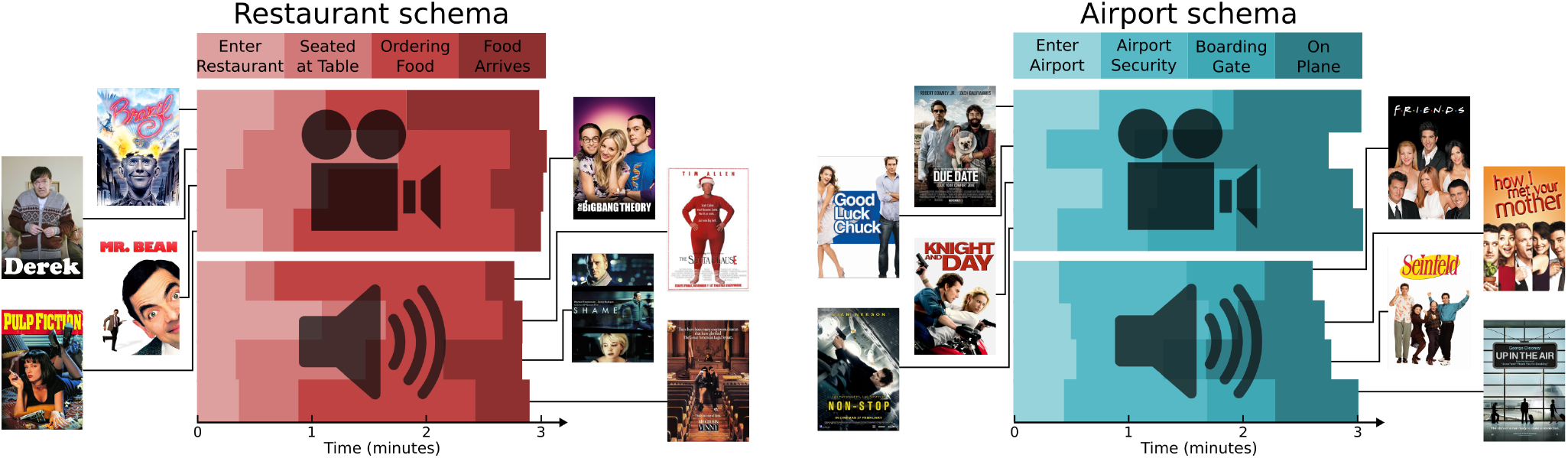
Experimental Stimuli. Subjects (N=31) were presented with 16 narrative stimuli, half audiovisual movies (clips from movies) and half audio narratives (based on movie scripts, all read by the same voice actor). The narratives varied widely, but conformed to one of two schemas: eating at a restaurant (entering the restaurant, being seated at a table, ordering food from a menu, and eating the food when it arrives) or going on a flight (entering the airport, going through security, waiting at the boarding gate, and taking a seat on the plane). The timing of these schematic events differed across stories, but all stories were approximately 3 minutes long.

To generate scrambled versions of the stimuli, each story was divided into its 4 schematic events, and then these clips were concatenated to create 16 new scrambled stimuli. Like the original stories, each of these scrambled stimuli contained all 4 schematic events from one schema and consisted of clips from a single modality (audio or audiovisual). Unlike the original stories, the schematic events in these clips were presented out of order, and were drawn from 4 different stories. 8 different “highly scrambled” permutations of 4 subevents (for which no two neighboring events were in the correct sequence, and at most 1 event was in the correct position in the sequence) were used for the 8 stories within each schema, as described in Table 1.

**Table 1:** Construction of scrambled stimuli. To test the effect of disrupting the overall temporal and narrative structure of the schematic events, we recombined the event clips of the same schema and modality into new, scrambled stimuli. Each of the 16 rows of the table indicates the four clips that made up each of the 16 scrambled stimuli, with the original event number of the clips indicated in parentheses.

| Schema | Modality | Clip 1 | Clip 2 | Clip 3 | Clip 4 |
| --- | --- | --- | --- | --- | --- |
| Restaurant | Audiovisual | Pulp Fiction (2) | Brazil (1) | Derek (4) | Mr. Bean (3) |
|  |  | Brazil (2) | Mr. Bean (4) | Pulp Fiction (3) | Derek (1) |
|  |  | Derek (3) | Mr. Bean (2) | Brazil (4) | Pulp Fiction (1) |
|  |  | Pulp Fiction (4) | Derek (2) | Mr. Bean (1) | Brazil (3) |
|  | Audio | Santa Clause (2) | Shame (4) | Big Bang (1) | Vinny (3) |
|  |  | Shame (3) | Vinny (1) | Santa Clause (4) | Big Bang (2) |
|  |  | Vinny (4) | Santa Clause (1) | Big Bang (3) | Shame (2) |
|  |  | Big Bang (4) | Santa Clause (3) | Vinny (2) | Shame (1) |
| Airport | Audiovisual | Non-stop (2) | Good Luck (4) | Knight (1) | Due Date (3) |
|  |  | Good Luck (3) | Due Date (1) | Non-stop (4) | Knight (2) |
|  |  | Due Date (4) | Non-stop (1) | Knight (3) | Good Luck (2) |
|  |  | Knight (4) | Non-stop (3) | Due Date (2) | Good Luck (1) |
|  | Audio | Up in the Air (2) | Friends (1) | How I Met (4) | Seinfeld (3) |
|  |  | Friends (2) | Seinfeld (4) | Up in the Air (3) | How I Met (1) |
|  |  | How I Met (3) | Seinfeld (2) | Friends (4) | Up in the Air (1) |
|  |  | Up in the Air (4) | How I Met (2) | Seinfeld (1) | Friends (3) |

All stimuli are publicly available at <https://figshare.com/articles/Event_Schema_Stimuli/5760306/3>.

### Scanning parameters and preprocessing

Data were collected on a 3T Siemens Prisma scanner with a 64-channel head/neck coil. Functional images were obtained with an interleaved multiband EPI sequence (TE = 39 ms, flip angle = 50°, multislice factor = 4, multi-slice shift = 3, FOV = 192 mm x 192 mm, partial Fourier = 6/8, 60 oblique axial slices) resulting in a voxel size of 2.0mm isotropic and a TR of 1.5s. The sequences used for the main experiment and scrambled control experiment were not identical (due to a sequence incompatibility caused by a scanner software upgrade) but had exactly the same parameters, except for a slightly different echo spacing (0.93 ms in the main experiment, 0.78ms in the control experiment). Whole-brain high-resolution (1.0 mm isotropic) T1-weighted structural mages were acquired with an MPRAGE sequence, and field maps were collected for dewarping (40 oblique axial slices, 3.0 mm isotropic).

Cortical surface extraction was performed on the anatomical image, using FreeSurfer 5.3. The Freesurfer epidewarp.fsl script was used to compute a voxel shift map to account for B0 distortion. The FreeSurfer Functional Analysis Stream (FsFast) was used to preprocess the fMRI data (alignment to the anatomy, motion and B0 correction, resampling to fsaverage6 cortical surface and subcortical MNI volume, 4mm smoothing). The resampled data (timecourses on the left and right hemispheres, and in the subcortical volume) were then read by a custom python script, which implemented the following preprocessing steps: removal of nuisance regressors (the 6 degrees of freedom motion correction estimates, and low-order legendre drift polynomials up to order (1 + duration/150) as in AFNI (Cox, 1996)), z-scoring each run to have zero mean and standard deviation of 1, and dividing the runs into the portions corresponding to each stimulus.

All subsequent analyses, described below, were carried out using custom python scripts and the Brain Imaging Analysis Kit <http://brainiak.org>. All ROI results can be fully reproduced online by using the published Code Ocean capsule <https://doi.org/10.24433/CO.a27d1d90-d227-4600-b876-051a801c7c20.v2>, which contains the python analysis code, preprocessed ROI data, and computational environment used for generating the results. The analysis code is also available on GitHub <https://github.com/cbaldassano/Schema-Perception>.

### Region of interest and searchlight definition

Based on prior work on the representation of high-level, cross-modal situation models (Zadbood, Chen, Leong, Norman, & Hasson, 2017), we focused our analysis primarily on regions within the default mode network. We derived these regions from an established network atlas on the fsaverage6 surface (Yeo et al., 2011), by selecting the networks from their 17-network parcellation (networks 15, 16, and 17) that made up the full default mode network (Buckner, Andrews-Hanna, & Schacter, 2008) and then merging parcels that were spatially contiguous. This yielded six regions of interest (ROIs): angular gyrus (1868 vertices), superior temporal sulcus (STS, 2118 vertices), superior frontal gyrus (SFG, 2461 vertices), medial prefontal cortex (mPFC, 2069 vertices), parahippocampal cortex (PHC, 882 vertices), and posterior medial cortex (PMC, 2495 vertices). Additionally, we used the freesurfer subcortical parcellation to extract the hippocampus as a region of interest (Hipp, 1289 voxels).

As a control region, we also defined an auditory cortex region based on correlations with the audio envelope of the stimuli. For every story (both movies and narratives), the root mean square amplitude of the audio signal was computed for every second of the stimulus. This was then convolved with a standard HRF (from AFNI, Cox, 1996) and downsampled to the temporal resolution of the fMRI signal. All audio envelopes were concatenated and then correlated with the concatenated timeseries from all stories at each surface vertex. The highest correlations were in the vicinity of Heschl’s gyrus, as expected, and an auditory cortex region was defined as all vertices with *r* > 0.27 in order to yield an ROI of comparable size to the other regions (1589 vertices).

Searchlight ROIs with a radius of approximately 15mm were generated by randomly sampling a center vertex, and then identifying all vertices within 11 steps of the center vertex along the surface mesh (since the vertex spacing of the fsaverage6 mesh is approximately 1.4mm, yielding a radius of 11*1.4mm≈15mm). Vertices without data (e.g. along the medial wall) were removed. Searchlights were randomly selected in this way until every vertex had been included in at least 3 searchlights.

For each ROI or each searchlight, data were aligned across subjects using the Shared Response Model (SRM) (P.-H. Chen et al., 2015). The goal of SRM is to project all subjects’ data into a common, lowdimensional feature space, such that corresponding timepoints from the same story are close together in this space. Given time by voxel data matrices *D_i_* from every subject, SRM finds a voxel by feature transformation matrix *T_i_* for every subject such that *D_i_* · *T_i_* ≈ *S*, where *S* is the feature timecourses shared across all subjects. We use data from all stories to estimate these transformation matrices, projecting all timecourses into a 100-dimensional space. Note that this projection will inflate the intersubject similarity for each story across subjects (since the transformations are chosen to maximize the similarity between corresponding timepoints), but will not artificially create similarity *between* stories, and does not use any information about the schema type of the stories.

### Experimental design and statistical analysis

After listening to a short unrelated audio clip to verify that the volume level was set correctly, subjects were presented with four stories in each of four experimental runs. Each run consisted of interleaved video and audio stories, with one story from each modality and schema in each run, and a randomized run order across subjects. Every story was preceded by a five second black screen followed by a five-second countdown video. The title of each story was displayed at the top of the screen throughout the story (the screen was otherwise black for the audio narratives). Subjects were informed that they would be asked to freely recall the stories after all sixteen had been presented (the recall data is not analyzed in this paper).

Statistics for all analyses were computed using nonparametric permutation and bootstrap techniques, as described in the sections below.

### Event pattern correlation analysis

First, the 31 subjects in the main experiment were randomly divided into two groups (of 15 and 16 subjects). For each story, four regressors were created to model the response to the four schematic events, along with an additional nuisance regressor to model the initial countdown video. These were created by taking the blocks of time corresponding to these five segments and then convolving with the hemodynamic response function from AFNI (Cox, 1996). A separate linear regression was performed to fit the average response of each group (in the 100-dimensional SRM space) using the regressors, resulting in a 100-dimensional pattern of coefficients for each event of each story in each group. For every pair of stories, the pattern vectors for each of their corresponding events were correlated across groups (event 1 from group 1 with event 1 from group 2, event 2 from group 1 with event 2 from group 2, etc., as shown in Figure 2a) and the four resulting correlations were averaged. This yielded a 16 by 16 matrix of across-group story event similarity. To ensure robustness, the whole process was repeated for 10 random splits of the 31 subjects, and the resulting similarity matrices were averaged across splits. For comparison, the same analysis was also performed without breaking the story into four segments (i.e., simply treating each story as a single event).

**Figure 2:**
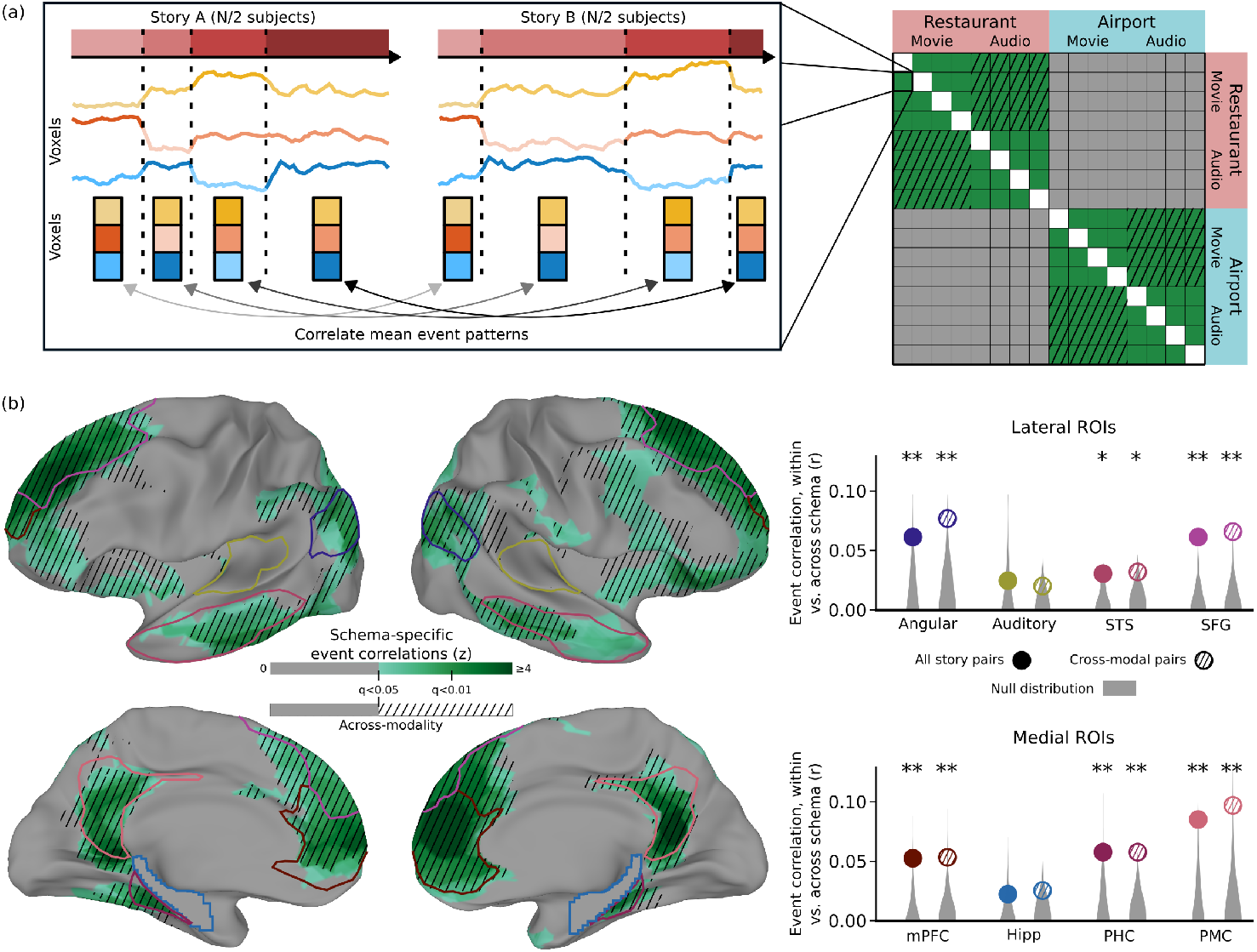
Event pattern correlations between stories. (a) For each pair of stories, we compute the mean pattern correlation (in a region of interest) between the 4 corresponding events. We then binned these correlations depending on whether the two stories were drawn from the same schema or different schemas. (b) We found that, throughout the default mode network, stories from the same schema showed significantly higher event similarity compared to stories from different schemas. This result was not driven solely by modality-specific stimulus features, since it appears even when considering only pairs of stories from different modalities (audiovisual movie and audio narrative). Note that auditory regions were not strongly related to high-level schemas, despite the presence of audio information in all stories. Surface map thresholded at FDR < 0.05; * p<0.05, ** p<0.01, by permutation test. Extended data are presented in Figure 2-1.

We then computed the average story similarity for pairs of different stories from the same schema versus pairs of different stories from different schemas. To determine if this difference was statistically significant, we randomly permuted the schema labels of the stories to generate a null distribution, and converted the true difference into a z value relative to this null distribution (yielding a corresponding p value from the survival function of the normal distribution). Additionally, we computed a more restrictive version in which only pairs of stories from opposite modalities (movie and audio narration) were considered, to ensure that schematic similarity was not driven by modality-specific features. A corresponding null distribution was generated similarly, by shuffling the schema labels of the stories (but keeping the modality labels intact). This analysis was applied within each of the ROIs and searchlights.

To explore the dimensionality of the schematic patterns, we re-ran the analysis after pre-processing the data with a range of different SRM dimensions, from 2 to 100. The resulting curve of z values versus dimensionality for each region was then smoothed with the LOWESS (Locally Weighted Scatterplot Smoothing) algorithm implemented in the statsmodels python package (using the default parameters).

To generate the final searchlight map, a z value was computed for each vertex as the average of the z values from all searchlights that included that vertex. The map of z values was then converted into map of q values using the same false discovery rate correction that is used in AFNI (Cox, 1996).

To visualize the spatial patterns, the participants were randomly split into two groups and a mean timecourse for each story was computed in each group. A regressor was created to model the response to all instances of each of the 8 schematic event types (across all stories), which were convolved with a hemodynamic response function and fit to each group’s data with a general linear model, as above (including the nuisance regressor to model the initial countdown video). Each resulting spatial pattern of coefficients was then z-scored and displayed on the cortical surface (masked to include only vertices that exhibited a significant schema effect in the searchlight analysis).

To determine the impact of scrambling the stimulus clips on the schema effect, we performed the same pattern correlation analysis described above, but in addition to correlating patterns between the two groups of subjects in the main (intact story) experiment, we also computed correlations between one of the intact groups and the scrambled control group. Note that SRM cannot be applied here (since the stimuli presented to the two groups are different) and so these analyses were conducted in the native vertex space. For assessing a statistically significant difference between the intact-intact and intact-scrambled correlations, the difference in schema effects in the two conditions (within-schema correlations minus across-schema correlations) was compared to a null distribution computed using the same permutation procedure described above.

### Schema classification analysis

An alternative analysis for detecting schematic structure was also performed, to determine if the schema type of novel stories could be decoded from fMRI data within an ROI, even when the novel stories have not been subdivided by hand into four events. The goal of this analysis was to use a labeled set of training stories to create a library of what each of the two schematic event sequences looked like on average (and measure how variable those patterns were across stories). Then, given held-out data from a new testing story, we attempted to divide that story’s timecourse into 4 events that looked like one of the 4-event sequences from our schematic event library. To perform this type of analysis in a principled way, we used a latent variable model in which every timepoint of the testing story belongs to some (unknown) event, and later timepoints must belong to later events than earlier timepoints.

Specifically, we used the Hidden Markov Model (HMM) variant introduced in Baldassano et al. (2017). This model assumes that activity patterns in a region proceed through a sequence of discrete latent states (starting with the first event and ending in the last event), and that within each event activity patterns are distributed in a Gaussian fashion around a mean event pattern. Given unlabeled timeseries data from a new story and a library of possible event sequences, we can infer the probability that each timepoint belongs to each of the schematic events.

Labels from 7 out of the 8 stories from each schema (the training set) were used to construct regressors for each of the 8 schematic event types, and then estimate a characteristic pattern for each event type using a general linear model (as in the visualization analysis above). We then measured the variance around these characteristic patterns within the corresponding events in the training stories. These mean and variance parameters were used to instantiate two HMMs, one loaded with the event patterns corresponding to the restaurant schema and the other with patterns from the airport schema. Each of the two HMMs was fit to each of the two held-out stories (one from each schema). This resulted in a (probabilistic) segmentation of the story into four events that matched the four corresponding events of the HMM as closely as possible. To evaluate the goodness of fit between the HMM and this story segmentation, we computed the four average patterns for the story timepoints assigned to each event, and compared them to the four characteristic HMM patterns by measuring the difference in mean correlation between corresponding and non-corresponding events. If the story is well-modeled by the HMM, the fitting procedure will be able to find four events (in the correct order) whose average spatial pattern is very similar to the four characteristic patterns in the HMM. Therefore, we attempted to classify which of the held-out stories came from which schema, based on which alignment of HMMs to stories provided a higher average match value.

The analysis described above was repeated for all possible choices of the two held-out stories. To obtain confidence intervals, we performed bootstrap sampling over subjects to produce 100 resampled datasets and ran the full analysis on each bootstrap sample. Note that, in order to keep data from different subjects independent (as is required for bootstrap sampling), we performed the fitting of the SRM after resampling.

To ensure that our classification performance was not biased by using training and testing data from the same run, we also ran an alternative version of this analysis in which the stories from 3 of the 4 runs were used as training data, and then the learned models were used to classify pairs of opposite-schema stories in the held-out run.

### Unsupervised alignment analysis

Finally, we sought to investigate whether the response to schematically-related narratives was consistent enough to automatically identify related events across stories, without relying on any human annotations. In contrast to the classification analysis above, fitting the HMM in this scenario requires not only inferring a (probabilistic) segmentation of each story into a series of shared events, but also estimating the shared event patterns themselves. This is accomplished by alternating between estimating the event patterns and inferring story segmentations until convergence, as described previously (Baldassano et al., 2017). The model therefore enforces that all stories must exhibit the same set of event patterns in the same order, but allows the durations of the events to vary across stories.

The number of latent events used (a hyperparameter of the HMM) was varied from 2 to 6 in steps of 1.

We then compared this model-predicted alignment to the hand-annotated event labels (defined as the maximum regressor of the event regressors described above). For any two stories, we found the set of pairs of timepoints (one from each story) that were predicted to be from the same event in the model (assigning each timepoint to its most probable event), and compared it to the set of timepoint pairs that had the same hand-annotated event label. We computed the intersection over union (the number of pairs in both sets divided by the number of pairs in either set) as a measure of how well the model correspondence matched the annotation correspondence. This similarity measure was averaged over all pairs of stories within a schema and results for both schemas were averaged. We generated a null distribution by replacing the model-defined event segmentation with a random segmentation, in which the event boundary timepoints were uniformly sampled from the set of all timepoints (without replacement), and performing the same analysis.

## Results

Our goal was to identify brain regions that represent the structure of real-life event schemas (e.g. eating at a restaurant) as they unfold over time (e.g. entering the restaurant, being seated, ordering, and eating food), irrespective of their low-level features, genres, characters, emotional content, and relative lengths of each of the constituent events. We therefore compared the activity patterns evoked by the schematic events of a story with those evoked by all other stories. We hypothesized that regions with schematic representations should show similarity between corresponding events of different stories with a shared schema but not stories with different schemas (Figure 2a).

Using both ROI analysis and a searchlight analysis, we found robust within-versus-between schema differences throughout the default mode network (Figure 2b), including the angular gyrus (p=0.009), STS (p=0.010), SFG (p < 0.001), mPFC (p < 0.001), PHC (p=0.005), and PMC (p=0.001). A borderline difference was also present in the hippocampus (p=0.066), and there was no significant difference in auditory cortex (p=0.137). The effect in auditory cortex was significantly smaller than that in angular gyrus (p=0.025), STS (p=0.043), mPFC (p=0.009), PHC (p=0.023), and PMC (p=0.002).

A more stringent requirement of schematic representation is that it should generalize across modalities. To test whether these effects were being driven solely by within-modality similarities (e.g. evoked by the visual objects or words associated with the schematic events), we repeated the same analysis but restricted our comparisons to pairs of story from different modalities (e.g. a spoken narration of a restaurant scene vs a movie clip of a restaurant scene). The pattern of significant effects was the same, including the angular gyrus (p=0.006), STS (p=0.030), SFG (p<0.001), mPFC (p=0.005), PHC (p=0.003), and PMC (p=0.002), with nonsignificant effects in the hippocampus (p=0.073) and auditory cortex (p=0.100). The searchlight analyses (performed both with all story pairs and with only across-modality story pairs) confirmed that the strongest effects occurred in these default mode ROIs, with weaker effects also evident in lateral prefrontal cortex and the insula.

To further explore the properties of the schematic spatial patterns driving these results, we repeated our analyses after projecting subjects’ data to a lower-dimensional shared space (Figure 2-1). This analysis revealed that schema effects were weaker in spaces with fewer than 10 dimensions, suggesting that the patterns we are measuring with fMRI reflect multiple distinct aspects of the stimuli and cannot be explained by a low-dimensional signal such as overall arousal or attention. Visualizing the characteristic spatial patterns for each of the 8 schematic events in two independent groups of subjects (Figure 3), we can observe that the spatial patterns are qualitatively both consistent across groups and distinct across schemas. The maps contain reliable spatial structure at a scale of approximately 1-2cm.

**Figure 3:**
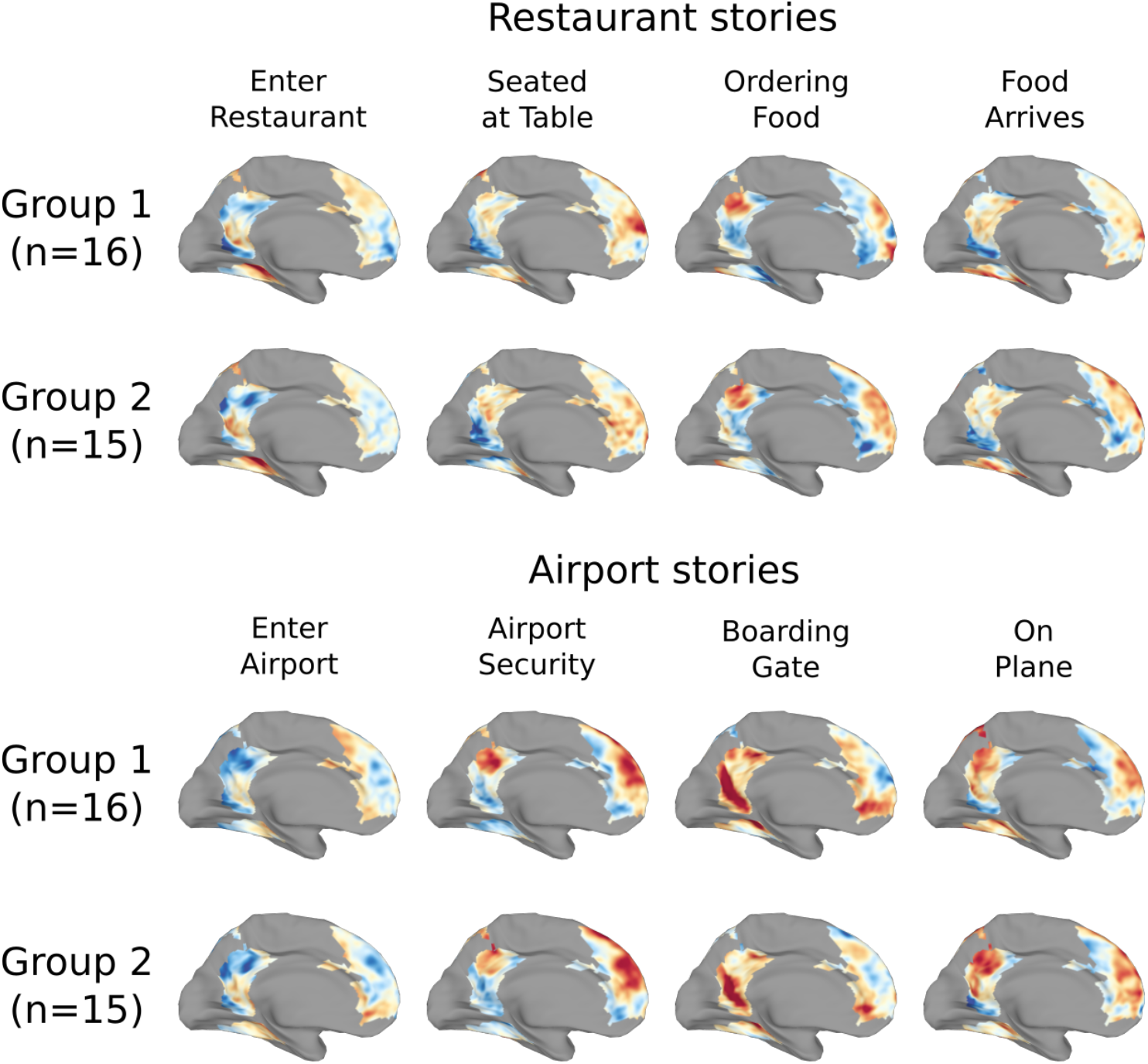
Visualization of schematic patterns. The spatial pattern associated with each of the schematic events was computed for two independent groups of subjects. Visualizing these patterns on the cortical surface (with warm colors indicating regions with above-average activity, and cool colors indicating regions with below-average activity, masked to show only those voxels that exhibited schema-related effects in Figure 2b) we observe that the spatial patterns are highly reliable across the split halves, and the patterns are visually distinct between the two schemas.

In the analyses above we divided each schema into four events (e.g. entering the restaurant, being seated, ordering, and eating food), and then measured the mean similarity of the patterns evoked by each event. An alternative analysis was conducted in which a single average pattern was used for each story (rather than a pattern for each event). This showed effects that were in the same direction (stronger within- than across-schema correlations), but substantially weaker (all p>0.12), suggesting that the representations in these regions are distinct for each of the stages of the schematic script.

These results indicate that default-mode regions represent schematic events in a way that generalizes across narratives. However, building schematic representations of specific events does not necessarily imply that these regions are tracking the full temporal script associated with restaurant or airport experiences. In order to determine whether these regions are sensitive to the longest-timescale structure across the narratives, we ran a control experiment with a separate group of subjects. Rather than viewing or listening to the intact stories, these subjects were presented with scrambled versions of the stories in which the schematic events were presented out of order and drawn from multiple stories within the same schema and modality (see Table 1). As before, we computed the spatial activity patterns evoked by the events of each story and compared them across stories to determine whether events of the same schematic type exhibited similar patterns. Note that the same stimulus events were being compared in both cases (e.g. the spatial pattern for the third event of *Brazil* in one group was correlated with the pattern for the third event of *Brazil* in the second group); however, the context in which those events had been presented to the subjects was different in the intact and scrambled groups. Therefore, brain regions that responded purely to the content of the current event should be unaffected by the scrambling, while brain regions sensitive to the overarching schematic context should show disrupted activity patterns and decreased schema effects.

As shown in Figure 4, we found that the scrambling significantly disrupted the schema effects in mPFC (p=0.028), and a post-hoc searchlight analysis within mPFC identified the ventral portion as the primary subregion impacted by the scrambling. Individual schematic events are therefore not sufficient to evoke strong schema-related activity patterns in mPFC without a coherent temporal script. Interestingly, none of the other regions showed a significant disruption of their schema effect with scrambling (all p>0.11), with some regions (such as angular gyrus and SFG) actually showing non-significant trends in the opposite direction (with disruption of a coherent narrative structure causing *more* schematic event representations).

**Figure 4:**
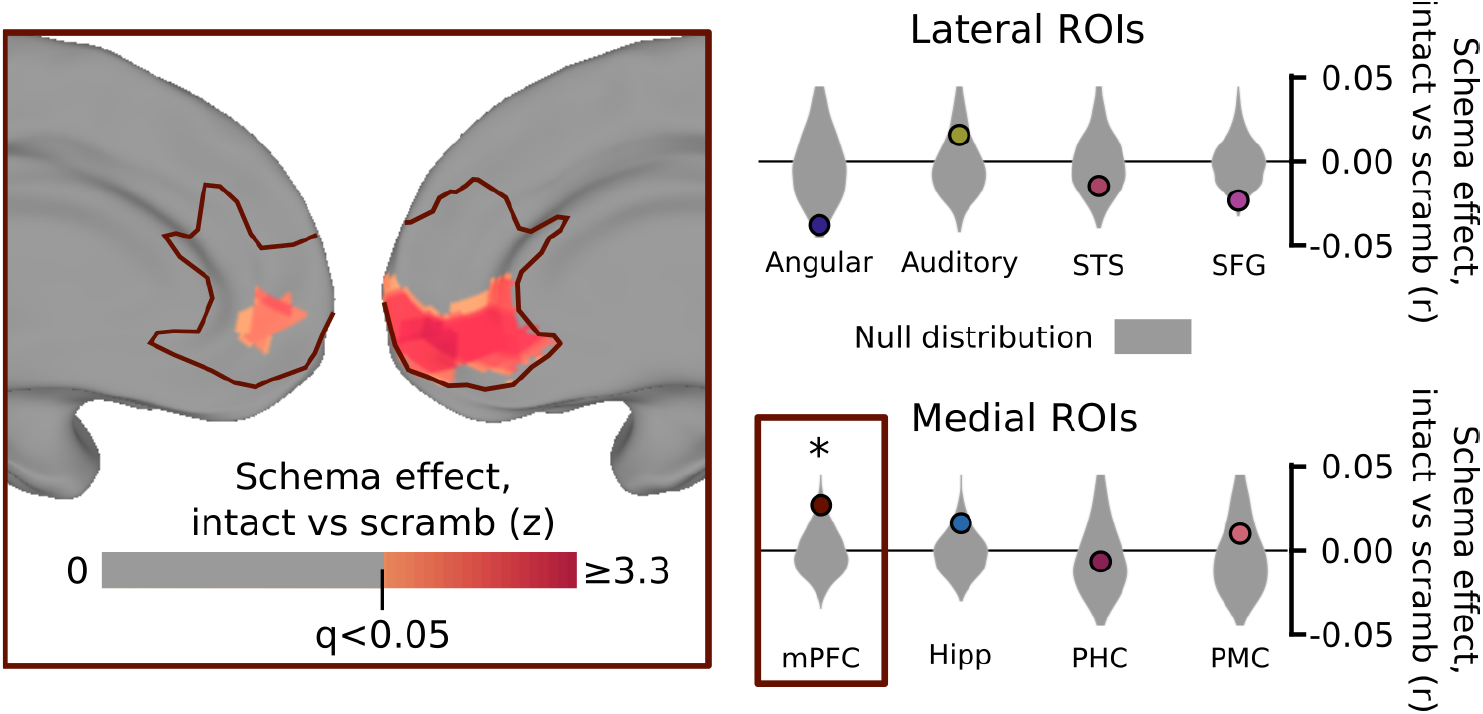
Effect of scrambling events on schematic correlations. A separate control group of 13 subjects was shown the same clips as in the main experiment, with the same schematically-blocked structure, but drawn randomly from different stories and in a random order. We compared the magnitude of the schema effect for story pair correlations between subjects in the main experiment (as in Figure 2) to the effect for correlations between subjects in the main experiment and the control experiment. Scrambling the event clips did not significantly disrupt schematic patterns in posterior regions of the cortex, but did significantly reduce the similarity between schematic events in mPFC, specifically in the ventral portion of mPFC (inset). This result shows that schematic patterns in mPFC are significantly enhanced by having intact, predictable script structure on the timescale of multiple minutes. Surface map thresholded at FDR < 0.05 within mPFC; * p<0.05 by permutation test.

We additionally performed a more stringent analysis to determine if a story’s schema (for intact stories in the main experiment) could be predicted from brain activity *without* labeling its temporal structure. Successfully predicting the schema of a held-out, unlabeled timeseries requires not just that the correctly-segmented event patterns are similar to the correct schema (as in the prior analysis), but also that there is *no* segmentation of the timeseries that yields event patterns similar to the incorrect schema. Using event annotations from 7 out of the 8 stories from each schema, we constructed characteristic activity patterns for each of the events from each schema. Using a Hidden Markov Model (Baldassano et al., 2017), we then attempted to segment the held-out stories into schematic events, with activity patterns matching the characteristic schema patterns from each schema. This allowed us to predict the schema type of the held-out, unlabeled story, based on which set of characteristic patterns best matched its evoked activity.

Even in this highly challenging classification task, we found significantly above-chance performance (Figure 5) in SFG (75% accuracy, p<0.001), mPFC (83% p<0.001), and PMC (69%, p<0.001). The other ROIs yielded lower, nonsignificant levels of performance (angular gyrus: 58%, p=0.142; auditory cortex: 57%, p=0.176; STS: 58%, p=0.209; hippocampus: 55%, p=0.305; PHC: 58%, p=0.149). Performing the analysis using an alternative cross-validation procedure (in which all stories from the same run were held out) yielded similar results (see Figure 5-1).

**Figure 5:**
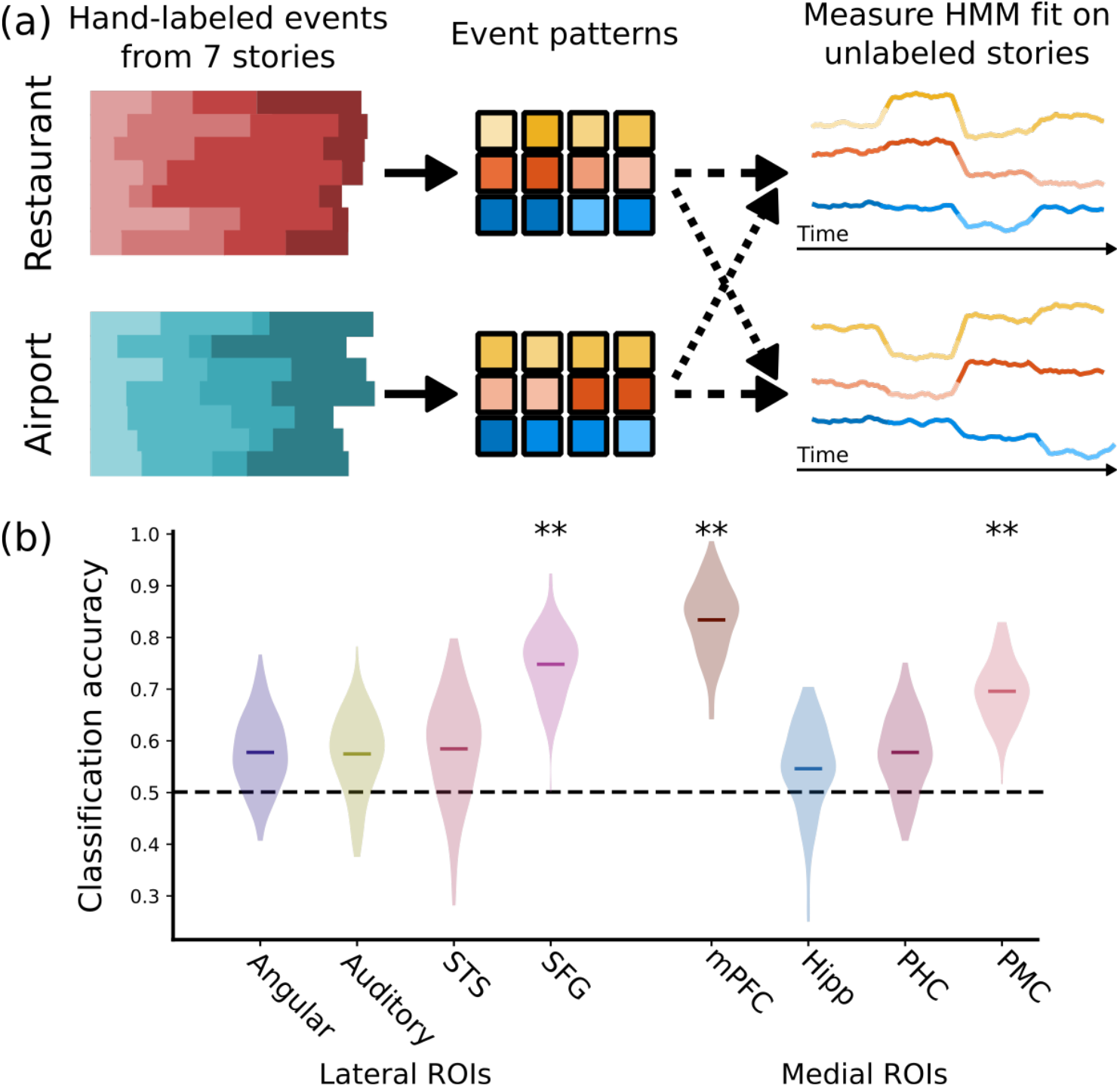
Schema classification of a new story. (a) Using the labeled event from 7 of the 8 stories (from each schema), we computed the average spatial activity pattern for each of the 4 events (from each schema), and used these as the latent event representations in two hidden markov models (HMMs). We then used the HMMs to find the best possible alignment of the held-out stories to each schema (without being given any event annotations), and predicted the schema of the held-out stories based on which of these alignments was better. (b) Superior frontal gyrus, medial prefrontal cortex, and posterior medial cortex all and medial prefrontal cortex showed classification rates well above chance, indicating that these regions exhibit robust schema-related patterns that can be identified in novel stories even without explicit event labels. ** p<0.01 by bootstrap test, shaded regions indicate bootstrap confidence distributions. Extended data are presented in Figure 5-1.

Finally, we employed data-driven methods to identify shared structure among stories within a schema, without the use of *any* prior knowledge of the schema’s temporal event structure (unlike the prior analysis, which used labeled annotations during training). Unlabeled timecourses from all eight stories within a schema were fit by an HMM, which sought to segment all stories into a sequence of events, such that the average activity patterns within each event were similar across stories (Figure 6a). We varied the number of latent events from 2 to 6, and measured how well the data-driven correspondence matched the hand-labeled schematic structure.

**Figure 6:**
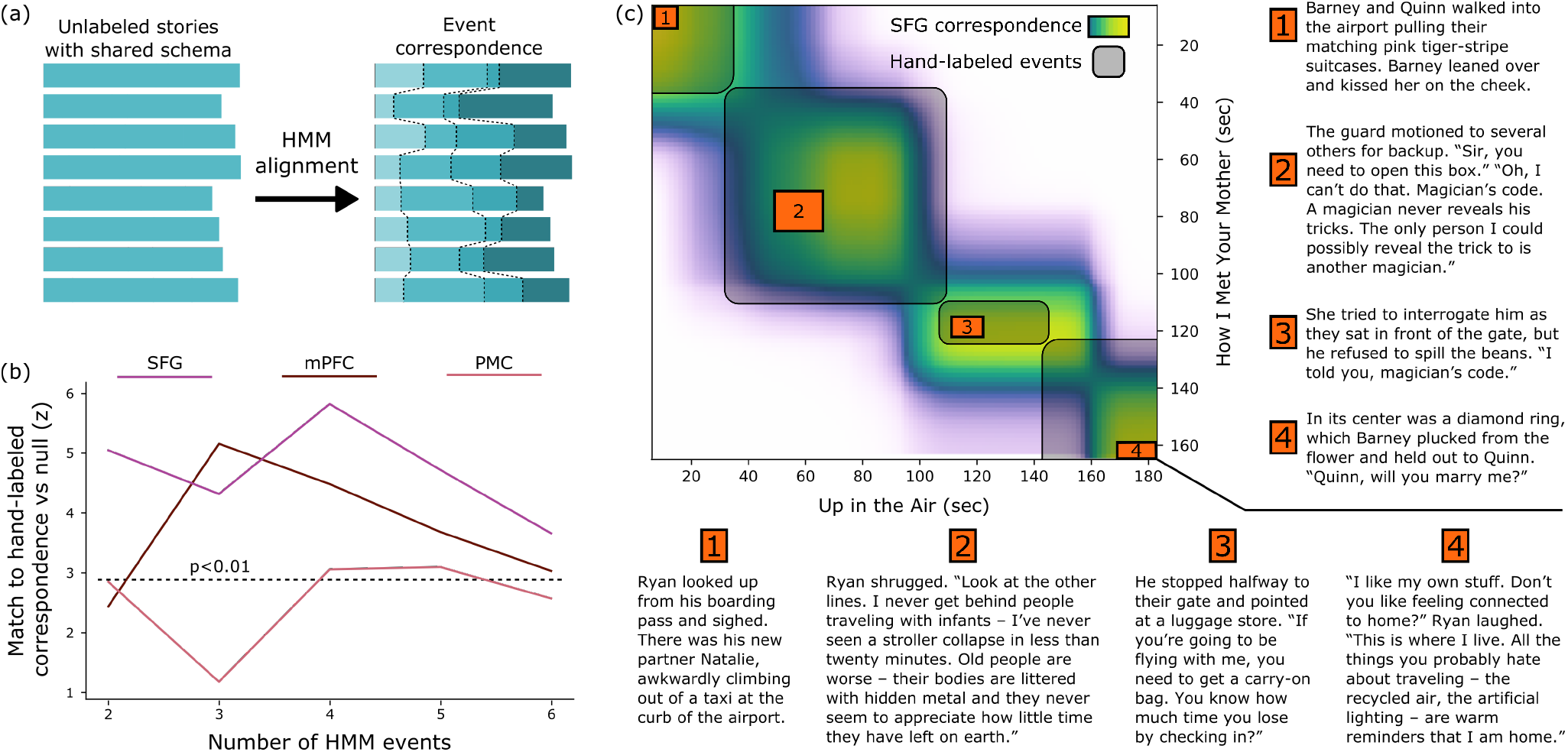
Unsupervised alignment of events across stories. (a) For each set of 8 stories within the same schema, we trained an HMM to identify a sequence of shared latent event patterns that was common across all the stories (without using any hand-labeled events). This correspondence can then be compared to the hand-labeled correspondence between stories. (b) We can recover the schematic structure shared across stories using only fMRI data, without event labeling, from all three ROIs showing strong schema representations in the previous analyses (SFG, mPFC, and PMC). This match is closest when the number of latent events in the model is similar to the true number of events (4). Dotted line indicates p<0.01 by permutation test, Bonferroni corrected for 5 tests. (c) For example, the data-driven correspondence between timepoints of “How I Met Your Mother” and “Up in the Air” (from SFG, with 4 events) captures the shared schematic events (entering the airport, going through security, waiting at the boarding gate, and taking a seat on the plane) despite large differences in the details of the narratives.

We found the SFG, mPFC, and PMC were all able to produce above-chance story alignments (Figure 6b), with the match peaking around 4 events (the number of schematic events the stimuli were constructed to have). Note that the correspondence was optimized simply to align the fMRI timeseries of different narratives and was not guaranteed to share any similarity with the hand-labeled annotations. As shown in an example alignment derived from the SFG data with four events (Figure 6c), the HMM is able to find correspondences across stories despite large variability in their specific content. These results indicate that schematic scripts are not only detectable in these regions (using the supervised analyses presented above) but are in fact a dominant organizing principle of their dynamic representations which can be discovered from an unsupervised alignment across stories.

## Discussion

Our results provide the first evidence that naturalistic perception activates dynamic information about the temporal structure of real-world scripts in mPFC, as well as a broader network including SFG and PMC. All three of these regions exhibited cross-subject, cross-modal shared representations of schematic events that could be used to classify the script type of held-out stories. Timecourses in these regions can be used to produce unsupervised alignments of stories with shared scripts, indicating that schematic representations are a primary driver of their ongoing representations. Additionally, the schematic representations in mPFC were disrupted when the events were presented outside the context of an intact temporal script, providing evidence that this region is involved in the selecting and maintaining script information on long timescales (on the order of multiple minutes). We also found weaker, less consistent evidence for schematic representations in the angular gyrus, hippocampus, STS, and PHC.

Prior work has shown that high-level regions including posterior medial cortex exhibit activity patterns that generalize across audio and video versions of the same story (Zadbood et al., 2017); this work extends this generalization to a further level of abstraction, showing similarities between *distinct stories* with very different content but a shared schematic skeleton. Our HMM-based classification approach (Baldassano et al., 2017) allows us to predict the schematic category of held-out stories with varying event timings purely from fMRI data, extending traditional decoding analyses into the temporal domain.

### Brain regions associated with schematic representations

There has been very little prior work investigating the neural mechanisms of real-world script-based perception. Work on simpler types of schematic representations have consistently implicated mPFC, based on encoding activity (Brod et al., 2016; van Kesteren et al., 2013, 2010), recall activity (Brod et al., 2015), gene expression (Tse et al., 2011), and lesion studies (Warren, Jones, Duff, & Tranel, 2014). Our results connect this large body of work with naturalistic scripts, suggesting that the same underlying neural mechanisms may be at play in both cases. Although the precise location of these mPFC results varies across studies, our peak effects fall squarely within an ROI (Yeo et al., 2011) belonging to the default mode network (Buckner et al., 2008), which has direct anatomical connections to posterior medial cortex (Greicius, Supekar, Menon, & Dougherty, 2009).

Schematic representations in mPFC are thought to interact with a network of posterior cortical regions, including posterior cingulate (van Buuren et al., 2014) and angular gyrus (Gilboa & Marlatte, 2017). Our results show strongly schematic responses in PMC and (less consistent) effects in angular gyrus, even in the absence of schema effects in mPFC, providing evidence that these regions intrinsically represent schematic information during perception. These regions are known to be sensitive to the structure of real-life events over relatively long timescales (minutes) (Baldassano et al., 2017; Hasson, Chen, & Honey, 2015), and are involved during a spoken replay of a narrative (J. Chen et al., 2017; Zadbood et al., 2017) suggesting that they encode and simulate sequences of actions in the world.

Finally, some studies have argued for schema formation within the hippocampus (McKenzie et al., 2014), while others have found that the hippocampus is most coupled to the cortex in the absence of a schema (van Kesteren et al., 2010). We found relatively weak effects in the hippocampus in all of our analyses. It is possible that extensively-learned scripts (such as the ones we used) are consolidated entirely into the cortex and are no longer mediated by the hippocampus (Norman & O’Reilly, 2003), or that hippocampal patterns would become schema-specific if subjects’ attention were explicitly directed to the schematic structure of the stories (Aly & Turk-Browne, 2016). Another possibility is that extensively-learned script representations are supported by highly sparse activity (possibly only present in a subregion of the hippocampus) that is not easily detectable using fMRI, or that requires specialized scanning sequences.

### Sensitivity to script structure in mPFC

Temporally scrambling natural stimuli has been a common approach to assess the sensitivity of brain regions to temporal structure at different timescales (Aly, Chen, Turk-Browne, & Hasson, 2018; Hasson et al., 2015; Lerner, Honey, Silbert, & Hasson, 2011; van Kesteren et al., 2010). In order to understand how the overall script context influenced event patterns, we scrambled our stimuli at the very coarse timescale of events (averaging 45 seconds long), and then compared the responses to the scrambled stimuli to those of the intact stimuli for pairs of stories from the same or different schemas. Our finding that only mPFC was sensitive to structure at this very long timescale is consistent with prior work showing that scrambling stories at the paragraph level (approximately 30 seconds) disrupted responses in mPFC, but that responses in other high-level regions such as PMC (after being temporally “unscrambled”) looked similar to those evoked by intact stories (Lerner et al., 2011). These results further support the view that mPFC is one of the few regions that track past context over long enough timescales to support the long-term temporal dependencies encoded in full naturalistic scripts.

### Building specific events from general scripts

In a previous study (Baldassano et al., 2017), we found that brain activity in posterior default mode regions jumps rapidly to a new, stable pattern at the start of a new event. This result raises a question: how can activity settle so quickly into a representation of the new event, when there has been very little time to accumulate information about the content of the event? Our results suggest a possible explanation of this phenomenon. If schematic script information is rapidly activated at the beginning of an event, and this information plays a critical role in setting the representations in these regions, then a substantial portion of the event representations can appear quickly at the beginning of an event. This proposal is similar to theories of visual perception, in which object associations are rapidly activated in mPFC and then used to influence representations in perceptual regions (Bar, 2007). This account is also consistent with our dimensionality analysis, which indicated that the event representations which were shared across subjects and drove the schema effects were relatively high-dimensional (spanning approximately a 10-dimensional space), suggesting that scripts can activate multiple distinct features of presented or inferred aspects of an event. In future work, approaches such as semantic modeling (Vodrahalli et al., 2017) may be able to disentangle the contribution of both schematic and specific content to the instantiation of event representations.

### Open questions

Although proposed models of schematic perception and memory have focused on mPFC and the medial temporal lobe (van Kesteren et al., 2012), our results raise the possibility that SFG and PMC could also play critical roles in representing the shared schematic structure of individual events. Schematic representations are present in these regions even in the absence of script representations in mPFC, suggesting that schematic representations in these regions may not be driven by top-down activation of scripts in mPFC but instead serve as the bottom-up building blocks for a complete narrative script. Further experiments will be required to identify the distinct functional contributions of these regions to perceptual and memory tasks, and to identify when and how they interact to produce schematic representations.

Another critical dimension is how representations in these regions develop over a lifetime (Brod, Werkle-Bergner, & Shing, 2013), since mPFC and its connections mature slowly throughout the first decade of life (Sowell, 2004; Supekar et al., 2010) and real-world event scripts can only be acquired after many exposures to events with shared structure. Both developmental studies and computational models of script learning could be used to understand how scripts develop in complexity, and whether perceptual representations become more script-based over time (Brod et al., 2013).

Finally, both this study and prior work on real-world scripts (Bower et al., 1979) have examined only a small fraction of the full library of scripts that adults have acquired over their lives, and have focused primarily on narrative perception. Additionally, individual scripts should not be studied only in isolation, since many real-world situations involve multiple simultaneously-active scripts (Schank & Abelson, 1977). New methodological approaches for studying perception in immersive real or virtual environments (Ladouce, Donaldson, Dudchenko, & Ietswaart, 2017) may allow us to sample more broadly from the ever-changing mix of regularities present in our daily lives.

## Extended Data

**Figure 2-1:**
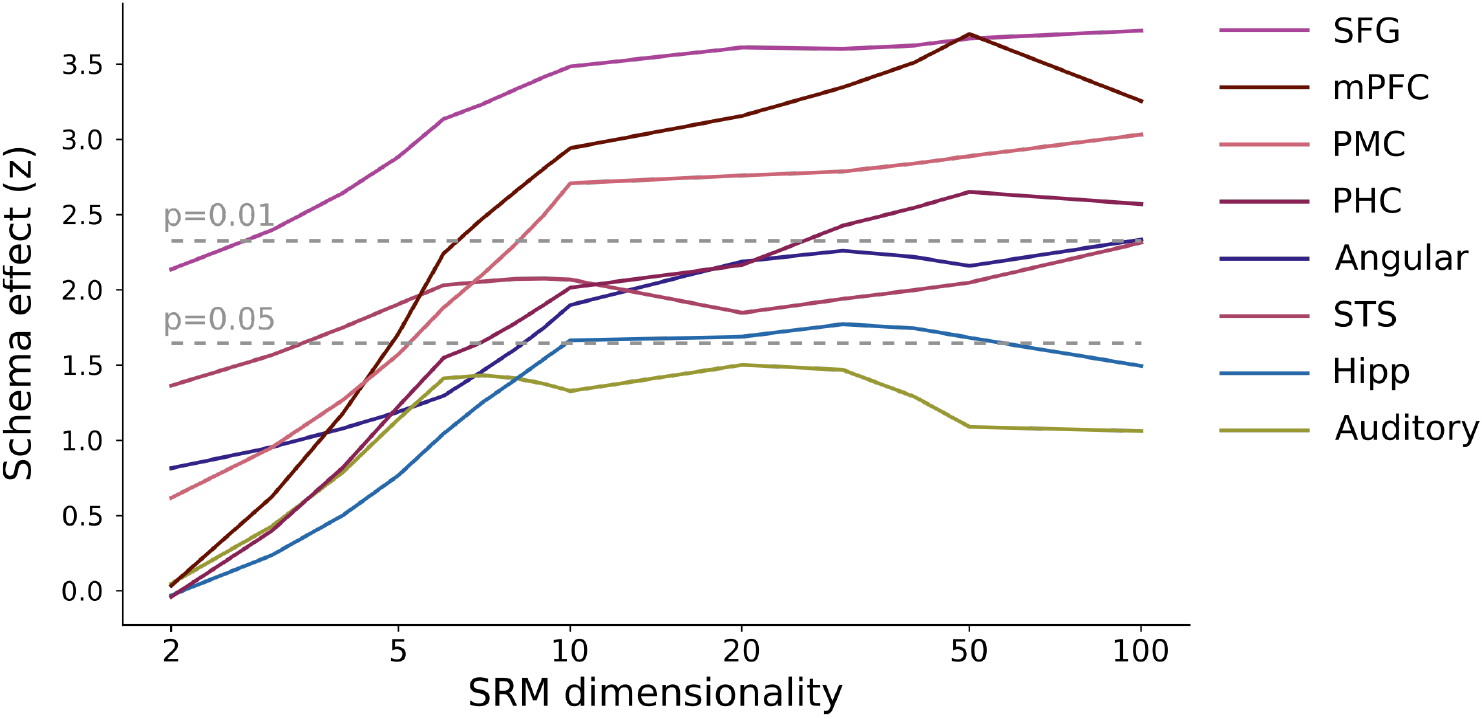
Dimensionality of event patterns. In order to estimate the underlying dimensionality of the spatial event patterns, the Shared Response Model (SRM) was used to project the group functional data into lower-dimensional spaces. With only two dimensions (the smallest possible dimensionality for which correlations can be computed) no regions show a schema effect significant at p<0.01 and only SFG shows an effect significant at p<0.05, indicating that a simple low-dimensional signal (such as global attentional modulation) is not sufficient to explain the results reported in Figure 2 (which used an SRM dimension of 100). Since the effects for all regions asymptote around 10 dimensions, we estimate that, across subjects, the fMRI signals of schematic event patterns span a roughly 10-dimensional space.

**Figure 5-1:**
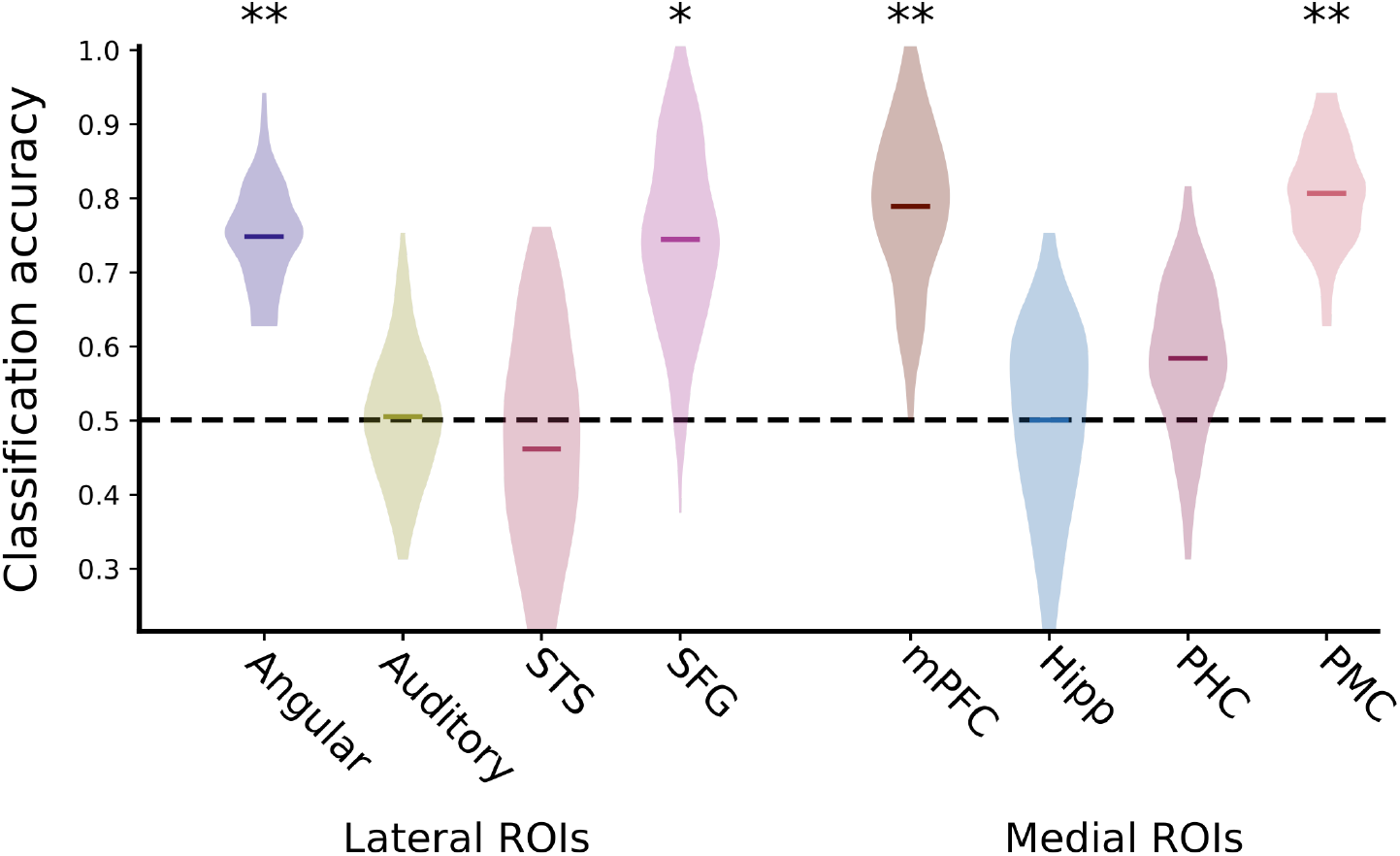
Schema classification, holding out stories from one run. Rather than classifying all pairs of opposite-schema stories, we can train the schema models on all but one run and then attempt to classify pairs of opposite-schema stories in the held-out run. As in Figure 5, classification accuracies above chance were observed in SFG (74% accuracy, p=0.024), mPFC (79%, p=0.004), and PMC (80%, p<0.001). Angular gyrus also exhibited above-chance accuracy (75%, p<0.001), while other regions were not significantly above chance (auditory cortex: 50%, p=0.483; STS: 45%, p=0.635; hippocampus: 50%, p=0.502; PHC:58%, p=0.209).

## Acknowledgements

We thank R. Masís-Obando for input on the Discussion, and the members of the Hasson and Norman labs for their comments and support. This work was supported by a grant from Intel Labs (CB) and The National Institutes of Health (R01 MH112357-01, UH and KAN).

## References

Aly, M., Chen, J., Turk-Browne, N. B., & Hasson, U. (2018). Learning Naturalistic Temporal Structure in the Posterior Medial Network. Journal of Cognitive Neuroscience, 25(9), 1–18. https://doi.org/10.1162/jocn_a_01308

Aly, M., & Turk-Browne, N. B. (2016). Attention promotes episodic encoding by stabilizing hippocampal representations. Proceedings of the National Academy of Sciences, 113(4), E420–E429. https://doi.org/10.1073/pnas.1518931113

Baldassano, C., Chen, J., Zadbood, A., Pillow, J. W., Hasson, U., & Norman, K. A. (2017). Discovering Event Structure in Continuous Narrative Perception and Memory. Neuron, 95(3). https://doi.org/10.1016/j.neuron.2017.06.041

Bar, M. (2007). The proactive brain: using analogies and associations to generate predictions. Trends in Cognitive Sciences, 11(7), 280–289. https://doi.org/10.1016/j.tics.2007.05.005

Bartlett, F. C. (1932). Remembering: A Study in Experimental and Social Psychology. Cambridge University Press. https://doi.org/10.1111/j.2044-8279.1933.tb02913.x

Bower, G. H., Black, J. B., & Turner, T. J. (1979). Scripts in memory for text. Cognitive Psychology, 11(2), 177–220. https://doi.org/10.1016/0010-0285(79)90009-4

Brod, G., Lindenberger, U., & Shing, Y. L. (2016). Neural activation patterns during retrieval of schema-related memories: Differences and commonalities between children and adults. Developmental Science, 1–16. https://doi.org/10.1111/desc.12475

Brod, G., Lindenberger, U., Werkle-Bergner, M., & Shing, Y. L. (2015). Differences in the neural signature of remembering schema-congruent and schema-incongruent events. NeuroImage, 117, 358–366. https://doi.org/10.1016/j.neuroimage.2015.05.086

Brod, G., Werkle-Bergner, M., & Shing, Y. L. (2013). The Influence of Prior Knowledge on Memory: A Developmental Cognitive Neuroscience Perspective. Frontiers in Behavioral Neuroscience, 7(October), 1–13. https://doi.org/10.3389/fnbeh.2013.00139

Buckner, R. L., Andrews-Hanna, J. R., & Schacter, D. L. (2008). The brain’s default network: anatomy, function, and relevance to disease. Annals of the New York Academy of Sciences, 1124, 1–38. https://doi.org/10.1196/annals.1440.011

Chen, J., Leong, Y. C., Honey, C. J., Yong, C. H., Norman, K. A., & Hasson, U. (2017). Shared memories reveal shared structure in neural activity across individuals. Nature Neuroscience, 20(1), 115–125. https://doi.org/10.1038/nn.4450

Chen, P.-H., Chen, J., Yeshurun, Y., Hasson, U., Haxby, J., & Ramadge, P. J. (2015). A Reduced-Dimension fMRI Shared Response Model. Neural Information Processing Systems Conference (NIPS), 460–468. Retrieved from http://papers.nips.cc/paper/5855-a-reduced-dimension-fmri-shared-response-model

Cox, R. W. (1996). AFNI: software for analysis and visualization of functional magnetic resonance neuroimages. Computers and Biomedical Research, an International Journal, 29(3), 162–73. Retrieved from http://www.ncbi.nlm.nih.gov/pubmed/8812068

Ghosh, V. E., Moscovitch, M., Melo Colella, B., & Gilboa, A. (2014). Schema Representation in Patients with Ventromedial PFC Lesions. Journal of Neuroscience, 34(36), 12057–12070. https://doi.org/10.1523/JNEUROSCI.0740-14.2014

Gilboa, A., & Marlatte, H. (2017). Neurobiology of Schemas and Schema-Mediated Memory. Trends in Cognitive Sciences, 21(8), 618–631. https://doi.org/10.1016/j.tics.2017.04.013

Greicius, M. D., Supekar, K., Menon, V., & Dougherty, R. F. (2009). Resting-State Functional Connectivity Reflects Structural Connectivity in the Default Mode Network. Cerebral Cortex, 19(1), 72–78. https://doi.org/10.1093/cercor/bhn059

Hasson, U., Chen, J., & Honey, C. J. (2015). Hierarchical process memory: Memory as an integral component of information processing. Trends in Cognitive Sciences, 19(6), 304–313. https://doi.org/10.1016/j.tics.2015.04.006

Ladouce, S., Donaldson, D. I., Dudchenko, P. A., & Ietswaart, M. (2017). Understanding Minds in Real-World Environments: Toward a Mobile Cognition Approach. Frontiers in Human Neuroscience, 10(January), 1–14. https://doi.org/10.3389/fnhum.2016.00694

Lerner, Y., Honey, C. J., Silbert, L. J., & Hasson, U. (2011). Topographic mapping of a hierarchy of temporal receptive windows using a narrated story. The Journal of Neuroscience: The Official Journal of the Society for Neuroscience, 31(8), 2906–15. https://doi.org/10.1523/JNEUROSCI.3684-10.2011

Mandler, J. M. (1984). Stories, scripts, and scenes: aspects of schema theory. Hillsdale, NJ: Erlbaum Associates.

McKenzie, S., Frank, A. J., Kinsky, N. R., Porter, B., Rivière, P. D., & Eichenbaum, H. (2014). Hippocampal representation of related and opposing memories develop within distinct, hierarchically organized neural schemas. Neuron, 83(1), 202–215. https://doi.org/10.1016/j.neuron.2014.05.019

Norman, K. A., & O’Reilly, R. C. (2003). Modeling hippocampal and neocortical contributions to recognition memory: A complementary-learning-systems approach. Psychological Review, 110(4), 611–646. https://doi.org/10.1037/0033-295X.110.4.611

Piaget, J. (1926). Language and thought of the child. New York: Harcourt, Brace & Co.

Schank, R. C., & Abelson, R. P. (1977). Scripts, plans, goals, and understanding: An inquiry into human knowledge structures. Lawrence Erlbaum Associates, Inc.

Sowell, E. R. (2004). Longitudinal Mapping of Cortical Thickness and Brain Growth in Normal Children. Journal of Neuroscience, 24(38), 8223–8231. https://doi.org/10.1523/JNEUROSCI.1798-04.2004

Stephens, G. J., Silbert, L. J., & Hasson, U. (2010). Speaker-listener neural coupling underlies successful communication. Proceedings of the National Academy of Sciences of the United States of America, 107(32), 14425–30. https://doi.org/10.1073/pnas.1008662107

Supekar, K., Uddin, L. Q., Prater, K., Amin, H., Greicius, M. D., & Menon, V. (2010). Development of functional and structural connectivity within the default mode network in young children. NeuroImage, 52(1), 290–301. https://doi.org/10.1016/j.neuroimage.2010.04.009

Tse, D., Langston, R. F., Kakeyama, M., Bethus, I., Spooner, P. A., Wood, E. R., … Morris, R. G. M. (2007). Schemas and Memory Consolidation. Science, 316(5821), 76–82. https://doi.org/10.1126/science.1135935

Tse, D., Takeuchi, T., Kakeyama, M., Kajii, Y., Okuno, H., Tohyama, C., … Morris, R. G. M. (2011). Schema-Dependent Gene Activation and Memory Encoding in Neocortex. Science, 333(6044), 891–895. https://doi.org/10.1126/science.1205274

van Buuren, M., Kroes, M. C. W., Wagner, I. C., Genzel, L., Morris, R. G. M., & Fernandez, G. (2014). Initial Investigation of the Effects of an Experimentally Learned Schema on Spatial Associative Memory in Humans. Journal of Neuroscience, 34(50), 16662–16670. https://doi.org/10.1523/JNEUROSCI.2365-14.2014

van Kesteren, M. T. R., Beul, S. F., Takashima, A., Henson, R. N., Ruiter, D. J., & Fernández, G. (2013). Differential roles for medial prefrontal and medial temporal cortices in schema-dependent encoding: From congruent to incongruent. Neuropsychologia, 51(12), 2352–2359. https://doi.org/10.1016/j.neuropsychologia.2013.05.027

van Kesteren, M. T. R., Fernández, G., Norris, D. G., & Hermans, E. J. (2010). Persistent schema-dependent hippocampal-neocortical connectivity during memory encoding and postencoding rest in humans. Proceedings of the National Academy of Sciences of the United States of America, 107(16), 7550–7555. https://doi.org/10.1073/pnas.0914892107

van Kesteren, M. T. R., Ruiter, D. J., Fernández, G., & Henson, R. N. (2012). How schema and novelty augment memory formation. Trends in Neurosciences, 35(4), 211–219. https://doi.org/10.1016/j.tins.2012.02.001

Vodrahalli, K., Chen, P.-H., Liang, Y., Baldassano, C., Chen, J., Yong, E., … Arora, S. (2017). Mapping between fMRI responses to movies and their natural language annotations. NeuroImage. https://doi.org/10.1016/j.neuroimage.2017.06.042

Warren, D. E., Jones, S. H., Duff, M. C., & Tranel, D. (2014). False Recall Is Reduced by Damage to the Ventromedial Prefrontal Cortex: Implications for Understanding the Neural Correlates of Schematic Memory. Journal of Neuroscience, 34(22), 7677–7682. https://doi.org/10.1523/JNEUROSCI.0119-14.2014

Yeo, B. T. T., Krienen, F. M., Sepulcre, J., Sabuncu, M. R., Lashkari, D., Hollinshead, M., … Buckner, R. L. (2011). The organization of the human cerebral cortex estimated by intrinsic functional connectivity. Journal of Neurophysiology, 106(3), 1125–65. https://doi.org/10.1152/jn.00338.2011

Zacks, J. M., Speer, N. K., Swallow, K. M., Braver, T. S., & Reynolds, J. R. (2007). Event perception: a mind-brain perspective. Psychological Bulletin, 133(2), 273–293. https://doi.org/10.1037/0033-2909.133.2.273

Zadbood, A., Chen, J., Leong, Y. C., Norman, K. A., & Hasson, U. (2017). How We Transmit Memories to Other Brains: Constructing Shared Neural Representations Via Communication. Cerebral Cortex, 27(10), 4988–5000. https://doi.org/10.1093/cercor/bhx202

